# High Cross Pollination Frequency in Rice (*Oryza sativa*) Landraces in Field Condition

**DOI:** 10.1101/2025.08.06.668876

**Authors:** Debal Deb, Debdulal Bhattacharya, Mahendra Nauri

## Abstract

Flower opening time (FOT) and the duration of florets remaining open (= flower exposure duration, FED) are crucially important for successful cross pollination between neighbouring rice plants. When the florets of the ovary parent (or pollen recipient) open even a few seconds after the closure of all the florets of the the pollen parent of a rice cultivar, the chances of cross pollination (CP) would obviously be zero. Conversely, as shown by our previous study, cross pollination frequency (CPF) would be very high (>80%) if the overlap of FEDs of the PP and OP is considerably long. In continuation of our previous study conducted in 2019, we subsequently conducted crossing experiments with 19 pairs of rice landraces during short-day flowering season and 8 pairs during long-day flowering season. The results show that with a long (>20 min) duration of FED overlap between OP and PP, the CPF tends to exceed 60%, often reaching 100%, although in some cases successful CP resulted in variable degrees of F1 sterility. These data confirm our previous report; and contradict the century-old ‘received wisdom’ based on a series of crossing experiments, that CP in rice rarely exceeds 2%, as widely published in all textbooks and official literature in rice biology. Our experimental evidence show the actuality of naturally high CPF in rice under circumstances of considerable FED overlap between parental cultivars, and indicates the necessity of further exploration into genetic interactions of sterile grain production and assortments of polygenic traits in the offspring from crosses between different varietal genotypes.

## 1. Introduction

The flower opening time (FOT) and the flower exposure duration (FED, measured as the period between flower opening time and closing time) are crucially important for the success of pollination and fertilization in rice plants. Although several studies over the past 115 years have reported the range of FOT and flower closing time (FCT) of different landraces (references in **Table 1**), none of them documented the FED of the pollen parent (PP) and ovary parent (OP) landraces used in any outcrossing experiments.

**Table 1:**
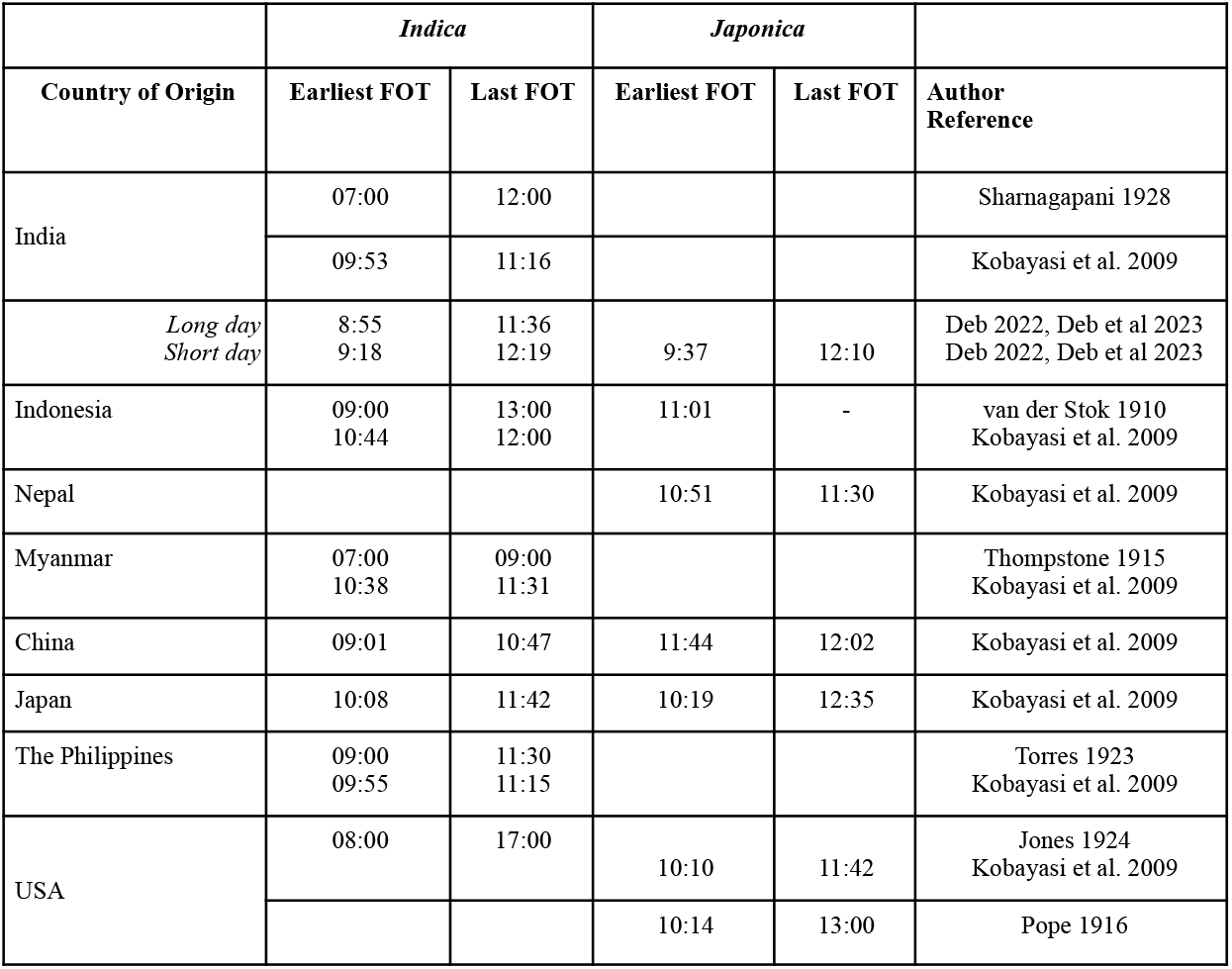
Published Records of Earliest and Latest FOT for Rice Cultivars from Different Countries.

The most crucial period for successful cross pollination is the pollen recipient flower’s exposure duration, calculated as the difference between the PP’s FCT and OP’s FOT, or alternatively, the OP’s FCT and PP’s FOT, which is denoted here as the ‘effective exposure duration’ (EED). The cross pollination frequency (CPF) is likely to be proportionately high only when EED > 0. In other words, if the flowers of the ovary parent (OP) close even a second before the FOT of the pollen parent (PP), the FED overlap between the two parents is zero, and consequently, there would be no cross pollination. Oblivious to this overlap of the crossing pair’s exposure duration (or EED as defined here), all previous experimental studies (Beachell et al. 1938; Coste 1969; da Silva et. 2005; Endo et al. 2009; Jodon 1959; Pope 1916; Reaño and Pham 1998; Robert et al. 1961; Sahadevan and Namboodiri 1963; Somaratne et al. 2012) as well as authentic reference books (Moldenhauer and Gibbonbs 1003; OGTR 2005) inform that cross pollination frequency (CPF) in cultivated rice (*Oryza sativa*) rarely exceeds 2%. This inference of very low CPF is also reported from experiments involving crosses between cultivated and wild rice (Langevin et al 1990; Song et al. 2003) as well as transgenic rice cultivars (Rong et al. 2007; Messegeuer et al. 2004). The only report of over 9% CPF, in one instance, came from an experiment by Bae et al. (2013), yet the overall CPF did not exceed 1% “in the field condition” (p. 168). We surmise that near-zero overlap of the FED between the PP and OP, unnoticed by the authors, might be the only factor resulting in extremely low CPF reported in those experiments, and therefore the received wisdom of extremely low CPF in cultivated rice is a severe underestimation of the actual CPF in real life conditions. Our previous study conducted in 2019 (Deb and Bhattacharya 2021) reported for the first time that CPF exceeded 80% in a pair of rice landraces, whose respective FEDs had significantly overlapped.

In continuation of our previous experiment in cross pollination conducted in 2019 (Deb and Bhattacharya 2021), we conducted similar experiments in subsequent years using 19 pairs of landraces during winter of 2020 and 8 pairs during summer of 2021. The range of flowering periods of the landraces in each pair were overlapping. We report here the results of the new sets of experiment to test our conjecture that the CPF in cultivated rice is usually very high unless the overlap between the FEDs of the crossing pairs is zero or near-zero.

## 2. Materials and Methods

### 2.1 Study site

The experiment was conducted at Basudha conservation farm (19°42′32.0′′N, 83°28′8.4′′E), located in Rayagada district, Odisha, India, during *aman* season (June–December) of the year 2020 and *boro* (Januiary – March) of 2021. The uniformly sandy loam soil of the farm was enriched with decomposed farm yard manure, farm-grown phosphate solubilizing microbial (PSM) inocula, and leaf mulching, and received no synthetic agrochemicals for the past 25 years.

### 2.2 Selection of landraces

The experiment was conducted with rice landraces in two seasons, first with the short-day (*aman*) season landraces, flowering in winter (October-November 2020); and subsequently, with the photoperiod-insensitive landraces flowering in long day (*boro*) season (April-May 2021).

Based on the previously recorded booting dates and unmistakable contrasting morphological characters of the landraces (Deb 2005; Bioversity International 2007), we selected a total of 26 pure lines of cultivars from Basudha’s repertoire of 1480 landraces, based on matching anthesis dates and FOT, and contrasting morphological characters (**Table 2**). The PP and the OP landraces were selected in order that they differed in at least 1 morphological trait. Among the 26 landraces, 16 were used as pollen parents and 19 as ovary parents for the crosses (**Suppl. Table S1**), comprising 19 parental pairs during *aman* and 8 pairs during *boro* season (**Suppl. Table S1**), totalling 27 pairs including 5 repeats.

**Table 2:** Countries of Origin and Heritable Morphological Characters of the Cultivars Used in Pair-wise Cross Pollination Experiments.

| Cultivar Code | Name | Origin | Pollen-<br>or Ovary<br>Parent | Flag Leaf<br>Attitude | Awn: Max<br>Length<br>(mm) | Hull<br>Colour | Bran<br>Colour | Kernel:<br>Notched<br>Belly % |
| --- | --- | --- | --- | --- | --- | --- | --- | --- |
| AA19 | Aurar | India | PP & OP | Droopy | 0 | Straw | Brown | 0 |
| B103 | Batapola El | Sri Lanka | PP & OP | Semi Erect | 0 | Straw | Light Brown | 0 |
| C34 | Chak ao Poreithon | India | PP | Erect | 14 | Black | Black | 0 |
| C44 | Choman | India | OP | Semi Erect | 0 | Brown<br>furrows | Brown | 0 |
| DD16 | Dude Bolta | India | OP | Erect | 25 | Gold<br>furrows | White | 0 |
| G25 | Gendi | India | OP | Horizontal | 0 | Purple<br>furrows | White | 0 |
| H01 | Haijam | India | PP & OP | Horizontal | 0 | Gold<br>furrows | White | 0 |
| H19 | Harir Jhinga | India | OP | Erect | 0 | Gold<br>furrows | White | 0 |
| H40 | Heen Dick Wee | Sri Lanka | OP | Erect | 0 | Brown<br>furrows | Light Brown | 0 |
| J33 | Juha | India | PP & OP | Horizontal | 0 | Gold<br>furrows | White | 0 |
| K131 | Kundapullan | India | PP | Droopy | 37 | Gold<br>furrows | White | >80% |
| K169 | Korgut | India | OP | Horizontal | 54 | Gold<br>furrows | Light Brown | 0 |
| L59 | Lankeswari | India | PP | Horizontal | 0 | Brown<br>furrows | White | 0 |
| M120 | Makhan | India | PP | Erect | 30 | Gold<br>furrows | White | 0 |
| M121 | Mala Gouri | India | PP | Semi Erect | 0 | Brown<br>(tawny) | White | 0 |
| M84 | Mavilon | Philippines | OP | Erect | 0 | Straw | Light Brown | 0 |
| M63 | Maw Thlen | India | OP | Erect | 0 | Straw | White | 0 |
| M91 | Mirlo | India | PP & OP | Erect | 0 | Brown<br>(tawny) | White | 0 |
| M94 | Muah | India | PP & OP | Semi Erect | 3 | Brown<br>(tawny) | Brown | 0 |
| N01 | Napa | India | PP & OP | Semi Erect | 2 | Straw | Light Brown | 0 |
| Q11 | Kharah | India | PP & OP | Semi Erect | 11 | Straw | White | 0 |
| S72 | Sahina | India | PP & OP | Erect | 0 | Straw | White | 0 |
| S87 | Samziuram Mei | India | OP | Horizontal | 6 | Straw | White | 0 |
| S20 | Sumitra | India | OP | Erect | 0 | Straw | White | 0 |
| T05 | Tiki | India | PP | Semi Erect | 0 | Purple<br>furrows | Black | 0 |
| V17 | Bhalu Dubraj | India | PP | Semi Erect | 62 | Black<br>furrows | White | >80% |

### 2.3 Selection of morphological characters

Each of the 26 landraces had already been assessed for 56 morphological characters to maintain genetic purity over the past 26 years (http://cintdis.org/basudha). We selected 5 visually distinctive morphological characters so that the PP and the OP of each pair differed in at least one of these 5 characters. Most of these characters were previously employed in crossing experiments by many authors (Beachell et al. 1938; da Silva et. 2005; Coste 1969; Endo et al. 2009; Reaño and Pham 1998; Robert et al. 1961; Sahadevan and Namboodiri 1963; Somaratne et al. 2012).

### 2.4 Experimental design

The experimental design of the crossing pairs was identical to our previous experiment ^[26]^. Experiments were set up on Basudha farm in two cultivation seasons – *Aman* (sown in June 2020, harvested in December 2020), and *Boro* (sown in February 2021, harvested in May 2021). During *Aman* 2020, 13 pairs of landraces were Selected, and 5 of the 13 pairs were replicated in two plots each. Thus the total number of plots for cross pollination (CP) was 18. A single seedling of a landrace, demarcated as the pollen recipient ovary parent (OP), was planted in the centre of each CP plot, around which 24 seedlings of a different landrace as the pollen parent (PP) were planted in a 5 × 5 matrix with 25 cm-spacing (**Fig. 1**, Plot A1 and Plot A2). The central position of the OP plant ensured the possible reception of pollen from all the surrounding PP plants, regardless of wind direction and speed. All seedlings were uniformly 14-day old when transplanted.

**Fig. 1:**
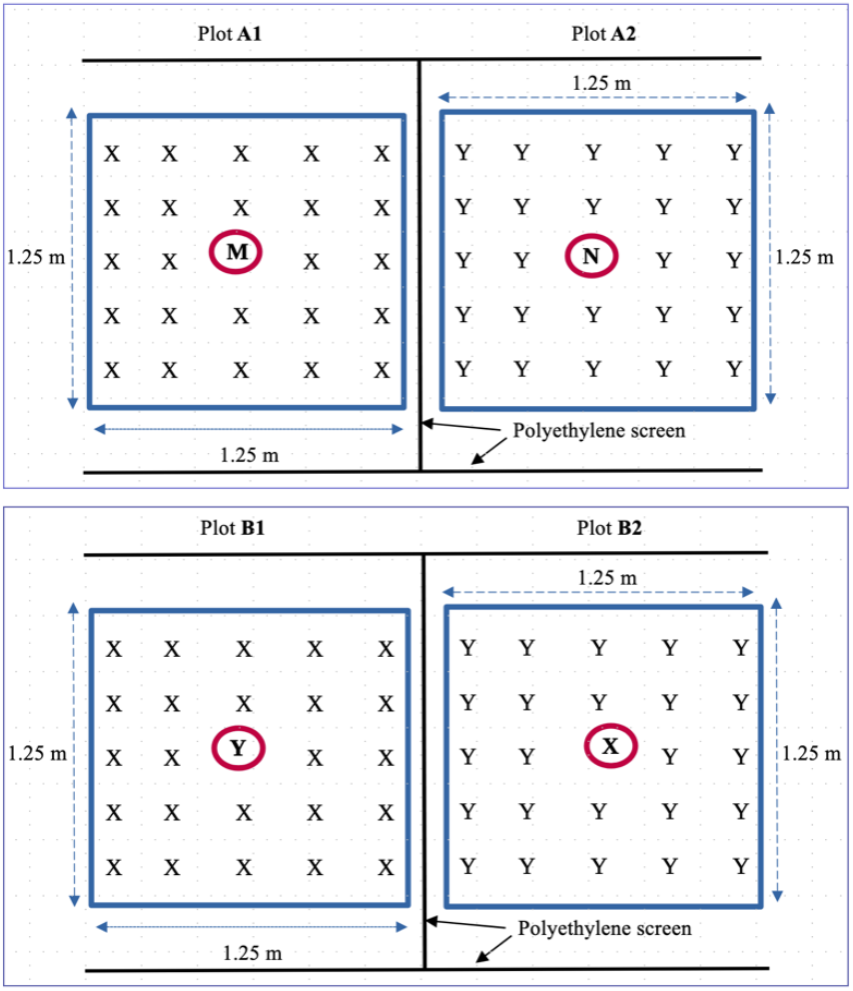
Planting Design of Two Experimental Plots with a Single Ovary Parent (OP, *in red circle*), Surrounded by 24 Pollen Parents (PP), Planted in a Density of 16 m^−2^. Plot A1 and Plot A2 Represent Different Pairs of PP and OP, planted during *Aman* season. Plot B1 and Plot B2 Represent Two Reciprocal Pairs (the OP in Plot B1 is the PP in Plot B2) Planted During *Boro* Season.

During *boro* season, 4 pairs of landraces were selected for reciprocal crossing, to examine whether the pollen flow from PP to OP remains effectively the same when the role of the parents are swapped. The arrangement of the each pair was reciprocally reversed in a corresponding neighbouring plot, such that each PP in a plot was planted as OP, and *vice versa*, in another plot (**Fig. 1**, Plot B1 and Plot B2). The total number of experimental plots with different pairs of these landraces, flowering during summer 2021, was 8.

At the booting stage, a thin polyethylene sheet was vertically placed as a barrier to pollen flux between the neighbouring plots. The height of the barrier was 1.5 m, always above the top panicle level in each plot.

### 2.5 Recording of FOT, FCT, FED and EED of the parental pairs

After the anthesis of each landrace, we recorded the first flower opening time (FOT) and the last floret closing time (FCT) in each varietal plot. The methods are given in detail in Deb et al. (2023). The longest duration of flowers remaining open (= flower opening duration, FED) was the interval between the FOT and the FCT of the last floret on a given varietal plot. As the transfer of pollen is possible only during the overlap of FED of OP and PP, we calculated the effective exposure duration (EED) of the OP florets, or the widest time window available for the OP’s florets receiving pollen. For any period during which both PP and OP florets remain open simultaneously, EED > 0, and EED = 0 otherwise.

### 2.6 Collection of F1 progeny seeds

After maturity of seeds, panicles of the F1 progeny were harvested from the single OP plant from each plot. The OP panicles of *Aman* season were harvested in January 2021, and the OP panicles from *Boro* season were harvested in April 2021. Isolation of each plot (**Fig. 1**) precluded seed contamination from any neighbouring plot. To preclude the possibility seed contamination, each mature panicle from the solitary 27 OP plants was enwrapped in a clean paper envelop, tagged, and harvested. Care was taken to clean our hands before and after enveloping each panicle. The harvested panicles were kept in the paper envelops until sowing in June 2021. Before sowing, the panicle density and sterility % were enumerated.

### 2.7 Cultivation of F1 plants

All the seeds from each panicle with labels bearing the parental pairs and corresponding plot numbers were sown in separate nursery beds on 20 June 2021. After germination, 14-to 16-d old F1 seedlings were transplanted in July 2021 in two rows in separate demarcated plots, where the soil had been prepared following standard organic method, with no synthetic agrochemical inputs.

### 2.8 Characterisation of the F1 offspring and CPF estimation

Every plant of the F1 progeny, germinated from viable F1 seeds, was visually examined for the selected morphological characters at maturity, namely, flag leaf angle, and the presence of awn. All panicles of the F1 plants were harvested after maturity in January 2022, and their seeds were assessed for hull (lemma and palea) colour, bran (pericarp) colour and the frequency of ‘notched belly’ trait (**Table 2**), to discern if any trait was inherited from the respective PP of the progeny. The scores of non-OP morphological traits from each F1 offspring were used to estimate the frequency inheritance of PP traits, whose data sheets are deposited in Harvard Dataverse repository (Deb 2025*b*).

### 2.9 Estimation of F1 sterility %

Inter-varietal cross pollination has been reported to yield variable proportions of F1 sterility (Engle 1969; Miller 1958, Zhang et al. 2022). However, natural population of most rice landraces show a non-zero grain sterility, reaching up to 12% (Deb 2005), which fluctuates in response to weather factors. For example, rainwash of the pollen may lead to a 100% grain sterility. However, in our experiments, no rainfall occurred during the flowering period of the landraces. Therefore we estimated the “control” grain sterility (Scontrol), from the data of the mean number of sterile grains per panicle in the population of each OP variety, routinely grown in Basudha rice farm field, and then estimated the *difference* between the grain sterility in the panicle of the solitary OP cultivar in each experimental plot (Sexp) and the sterility in the control plot (Scontrol) of the same cultivar, as the approximate frequency of F1 sterility due to cross pollination (Deb 2025*b*).

### 2.10 Assessment of gene flow frequency

The detection of at least one of the 5 contrasting characters of the PP in the F1 progeny would indicate the success of cross pollination. Since all these characters are a result of interactions of multiple genes, crosses between two landraces with contrasting traits is likely to produce offspring with variable quantifiable morphotypes. For a particular instance, at least one of the offspring of a long-awned PP crossed with an awnless OP may possess smaller awns on most grains. In this case, the presence of awn would be a clear evidence of inheritance of the PP trait, regardless of the exact length of the awn in the offspring. When both parental genotypes are identical for a specific character (e.g. same hull colour or the same bran colour), the F1 offspring will inherit the same trait from either parent. Therefore, we estimated only the non-OP trait inheritance alone. The cross pollination frequency (CPF) is a measure of the inheritance of at least 1 morphological character that is absent in OP, and was calculated as

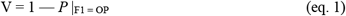

where V is the frequency of inheritance of non-OP characters in the viable progeny (germinated from non-sterile seeds), and *P* is the fraction of the viable F1 offspring inheriting all OP characters. However, considering that inter-varietal cross pollination often results in variable proportions of F1 sterility (Engle 1969; Miller 1958, Zhang et al. 2022), the equation 1 was modified to include the sterile seeds for estimation of the total transmission of PP genes to the offspring, including the sterile seeds:

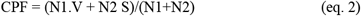

where N1 is the number of viable offspring, N2 is the number of sterile seeds, and S is the F1 sterility %, estimated as S = Sexp — Scontrol. When all F1 seeds are sterile, CPF = S. The CPF estimation procedure, derived from the data of non-OP morphological trait scores and F1 sterility, is shown in Harvard Dataverse repository (Deb 2025*b*).

## 3. Results

Using a total of 16 landraces as PP and 18 landraces PP, we planted selected pairs in a total of 26 experimental plots (including 5 replicates), of which 18 plots were established during *Aman* and 8 plots during *Boro* season (**Suppl. Table S1**). However, the entire progeny of one of these crosses (Q11 × S87 in plot A11) was lost due to insect pest attack before harvest. Therefore, we were unable to estimate the CPF from this pair. Furthermore, The F1 progeny of three crosses (plot A11, plot B2 and plot B4) produced 100% sterile grains (**Suppl. Table S2**).

The phenological details of flowering in the PP and OP landraces data are deposited in the Harvard Dataverse repository (Deb 2025*a*), and summarised here in **Fig. 2**, which shows that the EED of all pairs of landraces in this experiment ranged from 23 min (Plot A12a and Plot A12b) to 76 min (Plot A8). The mean of EED was 50.3 min during *Aman* season, and 56.8 min during *Boro* season. Each OP was expected to yield a progeny with a non-zero probability of inheritance of at least 1 trait from the corresponding PP.

**Fig. 2:**
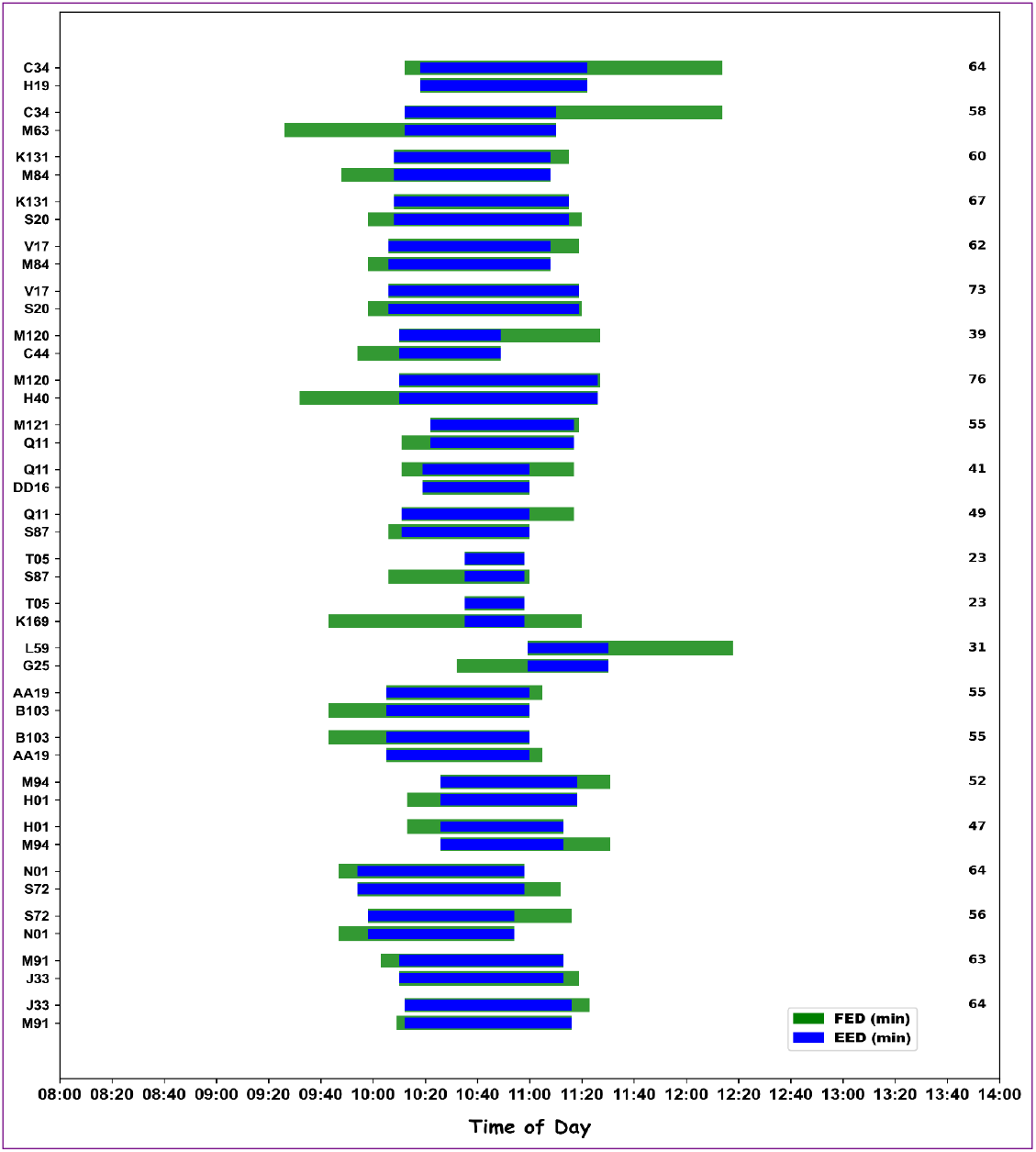
The Range of FED (green bar) Beginning from FOT and Ending at FCT of the PP (upper bar) and the OP (lower bar), with the Overlapping Period (blue) Indicating Effective Exposure Duration (EED) for Each Pair of Landraces. The column on the right margin shows the EED (in minutes) for each pair.

### 3.1 High F1 sterility in some crosses

Most of the crosses engendered variable proportions of F1 sterility. The crosses between the landraces in plot A13 (T05 × K169), plot B2 (B103 × AA19) and plot B4 (H01 × M94) produced 100% sterile grains. Some of the other crosses also produced a high proportion of sterile grains: the crosses between the landraces grown in plot A7a, plot A7b, plot A8, plot A11, plot A12 a, plot A12 b, plot A13, plot B6 and plot B7 produced >60% sterile grains borne on the respective OP’s panicle (**Fig. 3** and **Suppl. Table S2**). By contrast, plot A9 (M121 × Q11) produced no sterile seeds, and the crosses in plot A5 (V16 × M84), plot A6a and plot A6b (V17 × S20) respectively yielded 3.2%, 2% and 2.2% sterile seeds, in excess of the background rates (Scontrol) of 2%, 1.8% and 2.2%, respectively (**Fig. 3**). When the F1 sterility rates (estimated as S = [Sexp — Scontrol]; see Methods, **sec. 2.9**) were plotted against EED for all crosses, the slope of regression was significant at 99% confidence level (**Fig. 4**).

**Fig. 3:**
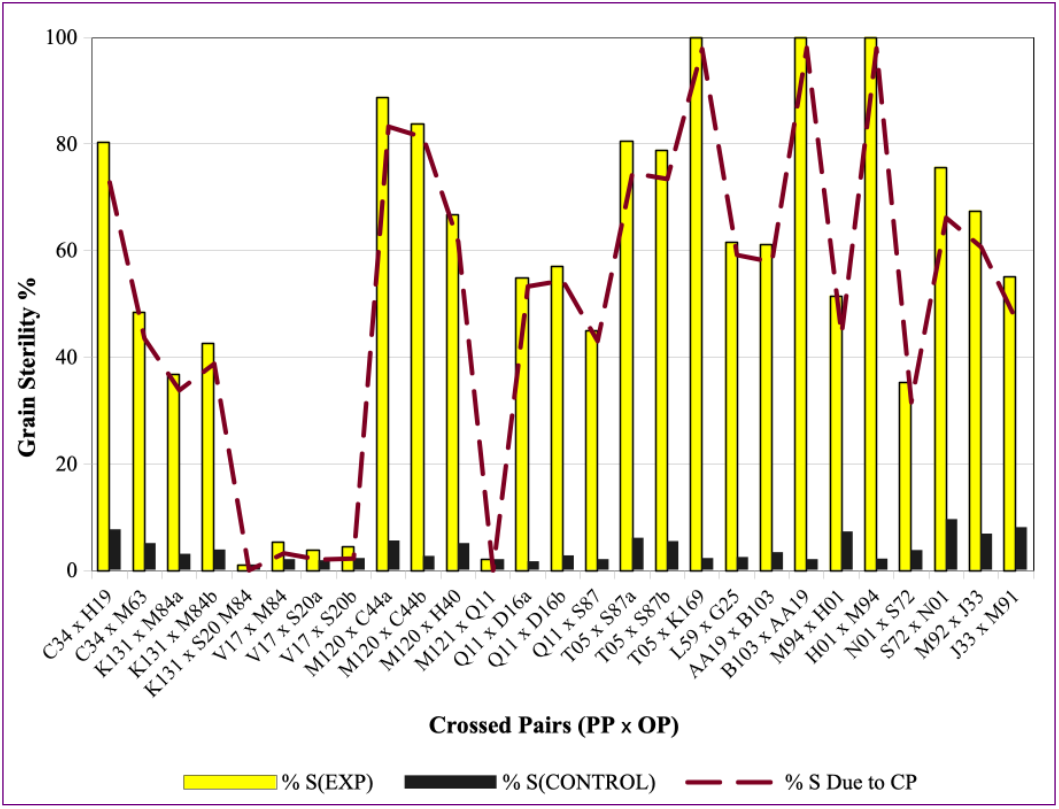
Estimation of Frequency of F1 Sterility Due to Cross Pollination as the Difference between Grain Sterility in the OP in CP Plots and Grain Sterility of the Isogenic Lines in Control Populations Grown in the Same Farm Field.

**Fig. 4:**
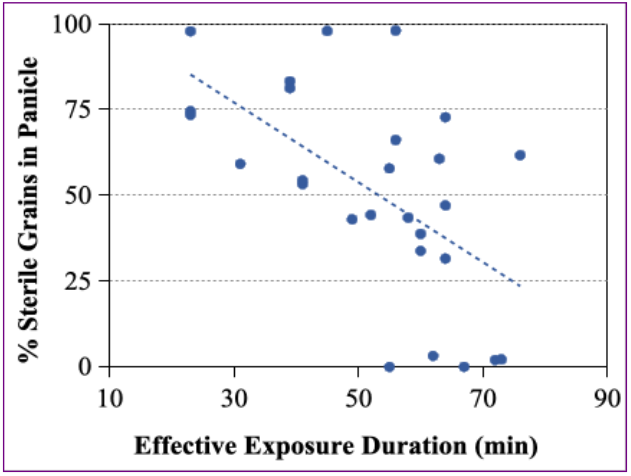
Regression of % Sterility of Grains in the Progeny of Crossed Parents against EED (min). Slope of regression = — 0.97, *df* = 25, *R*^2^ = 0.25, *t* = 2.74, *p <* 0.005.

### 3.2 High frequency of gene flow in all varietal pairs

The estimation of CPF was based on 5 unambiguous morphological markers, visually scored for each offspring. In the absence of any genetic contamination (assured by the experimental design), the visual scoring of at least one contrasting morphological markers in the offspring seems to be sufficient evidence of cross pollination. The frequency of inheritance of PP traits in each progeny from the experimental plots assigned to the corresponding parental pairs was precisely estimated using equation 2, and summarised in **Supplementary Table S2**). Excluding the progeny of a pair (Q11 × S87 in plot A11) destroyed by pest attack, 16 progeny lines were observed to have inherited > 60% of PP characters. No PP character was visible in the viable offspring in plot 1A (C34 × H19), implying zero inheritance in the viable offspring, although 80.4% of the total F1 seeds was sterile.

A sufficiently wide time window is logically conducive to successful cross pollination. Thus, the CPF is expected to be associated with the length of EED. When the proportion of inheritance of PP traits is plotted against EED, a definitive trend of CPF escalating with EED is evident, which can best described as a strong (*p* < 0.00001) 3rd degree polynomial relationship: % Inheritance = 157.6 — 0.001 EED^3^ + 0.19 EED^2^ — 6.57 EED (**Fig. 5**).

**Fig. 5:**
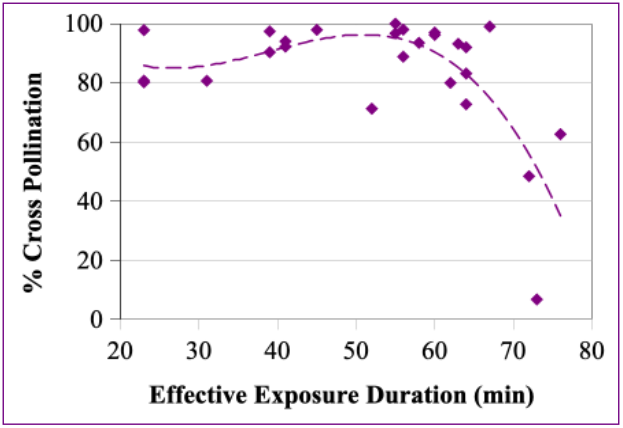
The Association of the Frequency of PP Trait Inheritance with EED of the OP, Best Described as a Polynomial Function (*df* = 25, *R*^2^ = 0.54, *t* = 5.26, *p* <0.00001).

## 4. Discussion and Conclusion

In the trivial case of zero- or near-zero overlap of FED between the PP and OP (EED ≈ 0), CPF ≈ 0; and this justifies the reports of extremely low CPF in most of the previous cross pollination experiments (Beachell et al. 1938; Coste 1969; da Silva et. 2005; Endo et al. 2009; Jodon 1959; Langevin et al 1990; Messegeuer et al. 2004; Reaño and Pham 1998; Robert et al. 1961; Rong et al. 2007; Sahadevan and Namboodiri 1963; Somaratne et al. 2012 Song et al. 2003). In our previous experiment (Deb and Bhattacharya 2021), one of the two parental pairs had an EED of ca. 10 min, and showed CPF = 0. In that same experiment, the FED overlap of 80 min was associated with CPF >80% — a result that is corroborated in the present study. Because pollen remains viable for no longer than 5 min after anther dehiscence, the long duration of FED overlap does not imply extension of pollen longevity. Rather, it implies that the ovary of an open floret remains receptive to viable pollen released from successively opening multiple florets borne on other panicles.

### 4.1 Alternative explanations for the expression of non-OP traits in F1

A major alternative source of non-OP characters in F1 progeny is seed contamination from surrounding rice plants during harvest. This possibility was precluded by (a) spatial confinement of each parental plots (**Fig.. 1**), (b) our seed harvesting protocol of enwrapping each OP panicle after maturity in a paper envelop, and (c) cleaning of hands before and after harvesting the OP panicles (see Methods, sec. 2.6).

The only other possibility of convergent segregation of characters in the OP ovaries is highly unlikely, because the same OP cultivars being separately cultivated on Basudha farm in the same season never produced a progeny bearing any of the non-OP characters. In particular, the divergence of any of the 5 distinctive characters selected in our study is not possible with polygenic interactions, unlikely to occur in any pure line genotype maintained on Basudha farm over 25 years.

The preclusion of seed contamination by our experimental design and procedure, and the extremely unlikely character segregation in the OP gametes indicate the veracity of a high observed frequency of transmission of PP traits in all the F1 progeny.

### 4.2 Two consequences of inter-varietal cross pollination

Cross pollination results in the transmission of the PP genes to the F1 progeny, but the inheritance of different morphological characters in rice does not follow simple Mendelian pattern. Because most of the rice characters are polygenic, different combinations of the genes from the pollen with the genes in the ovary produces an array of phenotypic descriptors. In addition, a large number of pleiotropic genes are linked to different biological pathways, governing heading date, grain length, plant height, flag leaf angle, and other characters (Li et al. 2023; Yao et al. 2018).

One of the 5 characters selected in this study is the ‘flag leaf angle’ phenotype, which is expressed in 4 categories, namely, erect, semi-erect, horizontal and droopy (Bioversity International 2007). The cross between M94 with semi-erect flag leaf (as PP) and H01 with horizontal flag leaf (as OP) produced a viable progeny with erect (15%), semi-erect (18%), and horizontal flag leaf (67%). Both the erect and semi-erect characters are non-OP characters, although the PP is a pure line with entirely semi-erect flag leaf. In the same pair, the pericarp colour of 55% of F1 seeds was light brown, although the rice kernels of both OP and PP had white bran. Likewise, the cross between L59 and G25 (plot A14), both parents characterised by horizontal flag leaf and white bran, produced 66% of the progeny with semi-erect flag leaf and 56% with light brown pericarp. Such (apparently unpredictable) departures from the parental characters in the offspring is because flag leaf angle and bran colour are complex phenotypes under polygenic control, with various associations of different genes on different chromosomes, each with multiple alleles (Jiang et al. 2022; Singh et al. 2024; Zhang et al. 2015). Low angles of flag leaf (erect and semi-erect) are partially dominant (Shen 1983; Zhang et al. 2015), and light brown bran colour is governed by a dominant gene in the absence of a complementary gene (Lee et al. 2018; Yang et al. 2022). The combinations of the multiple genes and their alleles in the zygote may result in the expression of an assortment of colour and flag leaf angles not exactly matching the character of either parent.

A second consequence of cross pollination is a range of seed sterility frequency, orchestrated by genetic interactions between the parental gametes F1 sterility, as reported in several studies (summarized in Miller 1958; Zhang et al. 2022). Low F1 fertility has been recorded mostly in *Oryza sativa* ssp. *indica* × *O. Sativa* ssp. *japonica* crosses, but also in some inter-varietal *indica* × *indica* crosses (Engle et al. 1969; Zhang et al. 2022), where the sterility genes determined by allelic interaction seem to be of wide occurrence. Different combinations of the *HSA1a* and *HSA1b* genes in *indica* rice are currently known to confer sterility in F1 and F2 hybrids (Kubo et al. 2016; Yang et al. 2022; Zhang et al. 2022).

The reciprocal crossing trials in 8 plots show that at least two pairs yielded a strikingly asymmetric result: In plot B1, the number of fertile grains of the progeny was 54, whereas the progeny of the reciprocal cross in Plot B2 produced no viable offspring (100% F1 sterility). Likewise, the cross between the pair grown in Plot B3 produced 66 offspring, whereas the reciprocal cross between the same landraces grown in Plot B4 produced only sterile grains (**Fig. 3**). Asymmetry in F1 sterility was also found in other crosses when a landrace is the PP, compared with the same landrace as OP. When Q11 was the PP in three crosses (plot 10a, plot 10b and plot 11), S ranged from 43% to 54%) whereas with Q11 as the OP (plot 9), S was 0% (**Fig. 3** and **Suppl. Table S2**).

Our observation of the two consequences of cross pollination in rice — F1 sterility (in excess of Scontrol) and inheritance of a combination of assorted phenotypes in the viable offspring — is summarised in a flowchart (**Fig. 6**), without allusion to biochemical and molecular genetic explanations, which are beyond the scope of this paper. This flowchart highlights the importance of FED overlap as well as the parental genotypes for cross pollination success.

**Fig. 6:**
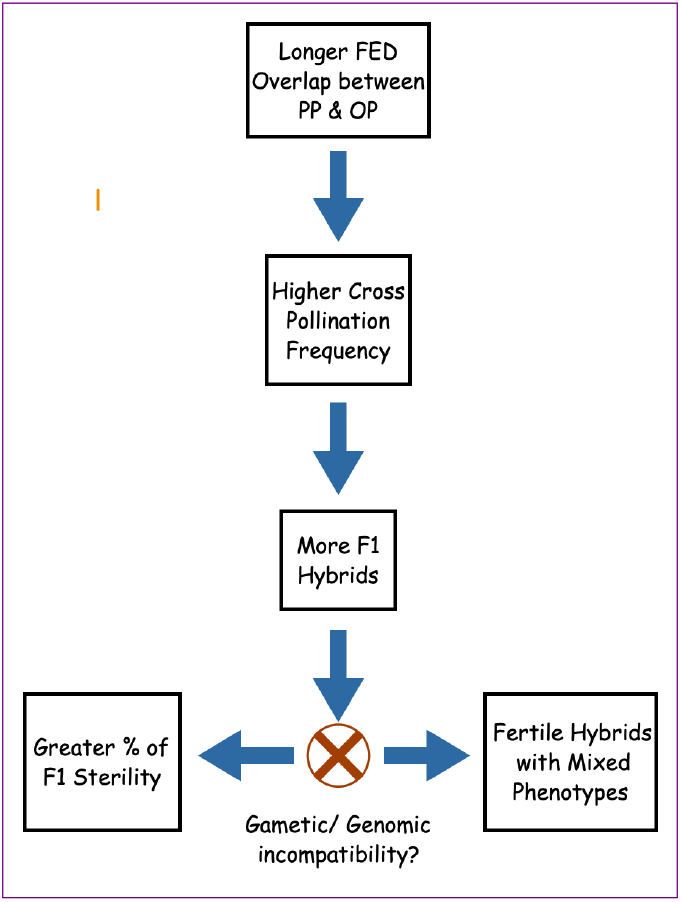
Flowchart Describing the Phenomenological Observations of Rice Phenology, Cross Pollination, and its Consequences in Field Condition.

### 4.3 The implication of F1 sterility in CPF estimation

Because grain sterility is a consequence of cross pollination itself, all the flowers of the solitary OP plants in both Plot B2 and Plot B4 can safely be surmised to have been cross pollinated (CPF =100). Taken together, our study presents a convincing empirical evidence of considerably high CPF between *indica* landraces, as long as the overlap of FED between PP and OP flowers is of sufficiently long (>20 min) duration. The high frequency of CPF and gene flow reported here is in sharp contradiction to the prevalent knowledge of ‘less than 2% CPF’ presented in all biology text books and official rice biology reports. The high frequency of inheritance of PP traits by the progeny documented in this study establishes the factual possibility of high CPF between landraces in field conditions, corroborating our earlier finding (Deb and Bhattacharya 2021).

The results of this study seem to be conclusive of the actuality of naturally high CPF in rice under circumstances of considerable FED overlap between parental cultivars, and indicates the necessity of further exploration into the biochemical pathways and genetic interactions of sterile grain production and assortments of polygenic traits in the offspring from crosses between different varietal genotypes. The latter investigation may consolidate our contention that contrary to the the extremely low CPF that is generally accepted in the mainstream rice literature over a century, CPF in field conditions is much higher, depending on FED overlap between neighbouring cultivars. These findings are of profound importance in successful hybridization efforts (by selecting cultivars with wider EED); and conversely, in efforts to maintain genetic purity of rice landraces (by planting multiple landraces next to one another with non-overlapping EEDs). The rice breeders’ and farmers’ concern of unintended genetic contamination of their crops from transgenic (such as herbicide tolerant) rice pollen flow can also be addressed by de-synchronising the FOT and FED of the OP from that of the transgenic PP cultivars.

## Acknowledgements

We are immensely grateful to Prof. N V Joshi of Indian Institute of Science, Bangalore for his meticulous scrutiny of data and critical comments on earlier drafts of the mss. We are indebted to Sri Avik Saha for all logistic support to our work on Basudha farm and laboratory.

## Data availability

In addition to the data provided in Supplementary Tables, phenological details of 26 rice landraces are available on Harvard Dataverse (Deb 2025*a*). The details of calculation of the frequency of hybrid sterility and of the frequency of transmission of PP into F1 offspring (**Suppl. Table S2**) are available on Harvard repository (Deb 2025*b*). Any other data will be freely available on reasonable request.

## Author Contributions

Conceptualization: DD; Design & Methodology: DD; Field investigation, Data collection, Data entry: DB, MN; Data curation: DD; Formal analysis: DD; Writing – original draft: DD; Writing – review & editing: DD, DB.

## Statements and Declarations Funding

No funding was received for conducting this study.

## Competing interests

The authors have no relevant financial or non-financial interests to disclose.

## Notes

https://doi.org/10.7910/DVN/TZCNYR

https://doi.org/10.7910/DVN/QZIV0H

### Competing Interest Statement

The authors have declared no competing interest.

### Summary of Updates

A section has been added to the older version to discuss the obviation of alternative explanations (such as seed contamination) of high CPF reported in this study. Also rectified an error in a supplemental table.

