## Supplementary Table S1 for "High Cross Pollination Frequency in Rice (*Oryza sativa*) Landraces in Field Condition"

**Supplementary Material**

**Table S1**: Experimental Design Matrix Showing Crosses between 16 Landraces as Pollen Parents and 19 Landraces as Ovary Parent. Cell entries indicate Plot Numbers allotted to discrete pairs.

| | **Aman 2020** | | | | | | | | **Boro 2021** | | | | | | | |
|---|---|---|---|---|---|---|---|---|---|---|---|---|---|---|---|---|
| ***PP* ▶**<br>***OP* ▼** | **C34** | **K131** | **L59** | **M120** | **M121** | **Q11** | **T05** | **V17** | **AA19** | **B103** | **H01** | **J33** | **M91** | **M94** | **N01** | **S72** |
| **H19** | A1 | | | | | | | | | | | | | | | |
| **M63** | A2 | | | | | | | | | | | | | | | |
| **M84** | | A3a<br>A3b | | | | | | A5 | | | | | | | | |
| **S20** | | A4 | | | | | | A6a<br>A6b | | | | | | | | |
| **C44** | | | | A7a<br>A7b | | | | | | | | | | | | |
| **H40** | | | | A8 | | | | | | | | | | | | |
| **Q11** | | | | | A9 | | | | | | | | | | | |
| **DD16** | | | | | | A10a<br>A10b | | | | | | | | | | |
| **S87** | | | | | | A11<br>$ | A12a<br>A12b | | | | | | | | | |
| **K169** | | | | | | | A13<br>$ | | | | | | | | | |
| **G25** | | | A14 | | | | | | | | | | | | | |
| **B103** | | | | | | | | | B1 | | | | | | | |
| **AA19** | | | | | | | | | | B2<br>$ | | | | | | |
| **M94** | | | | | | | | | | | B4<br>$ | | | | | |
| **M91** | | | | | | | | | | | | B8 | | | | |
| **J33** | | | | | | | | | | | | | B7 | | | |
| **H01** | | | | | | | | | | | | | | B3 | | |
| **S72** | | | | | | | | | | | | | | | B5 | |
| **N01** | | | | | | | | | | | | | | | | B6 |

**a** and **b** denote replications of the same pair
Plots marked by a $ sign denote lost/ discarded morphological data of the F1 progeny (see text).
