## Supplementary Table S2 for "High Cross Pollination Frequency in Rice (*Oryza sativa*) Landraces in Field Condition"

**Table S2**. Estimation of % Inheritance of Characters in F1 Not Matching OP characters in Crosses between 25 Pairs of Landraces.

| Plot No. | Pollen Parent | Ovary Parent | F1 Panicle Density | Hybrid sterility % | Total Grain Sterility % | No. of Fertile Seeds | F1 Seeds Lost to Pest | No. of Viable F1 Offspring | Non-OP Characters in F1 (%) | | | | | % Viable Progeny with Non-OP traits * | CPF # |
|---|---|---|---|---|---|---|---|---|---|---|---|---|---|---|---|
| | | | | | | | | | Flag Leaf Angle | Awn | Hull Colour | Bran Colour | Notched Belly | | |
| **A1** | C34 | H19 | 112 | 72.8 | 80.4 | 22 | | 22 | ? | 0 | 0 | 0 | ? | 0 | 72.8 |
| **A2** | C34 | M63 | 132 | 43.5 | 48.5 | 68 | | 68 | 97.10 | 72.06 | 0 | 0 | ? | 97.1 | 93.5 |
| **A3a** | K131 | M84a | 193 | 33.8 | 36.8 | 122 | | 122 | 0 | 67.21 | 100 | | 35.25 | 100 | 97 |
| **A3b** | K131 | M84b | 188 | 38.8 | 42.6 | 108 | | 108 | 0 | 74.07 | 93.52 | | 35.19 | 100 | 96.2 |
| **A4** | K131 | S20 | 198 | 0.0 | 1.0 | 196 | | 196 | 13.27 | 0.51 | 100 | ? | 16.33 | 100 | 99 |
| **A5** | V17 | M84 | 135 | 3.2 | 5.2 | 128 | | 128 | 0 | 81.00 | 0 | 0 | 32.81 | 81.2 | 80.2 |
| **A6a** | V17 | S20a | 228 | 0.4 | 2.2 | 223 | | 223 | 8.52 | 0 | 0 | ? | 47.09 | 47.5 | 46.9 |
| **A6b** | V17 | S20b | 220 | 2.3 | 4.5 | 210 | 154 | 59 | 16.95 | 0 | 0 | ? | 1.69 | 16.9 | 6.9 |
| **A7a** | M120 | C44a | 142 | 83.2 | 88.7 | 16 | | 16 | 62.50 | 0 | 13 | 0 | ? | 66.7 | 90.7 |
| **A7b** | M120 | C44b | 136 | 81.2 | 83.8 | 22 | | 22 | 45.45 | 0 | 0 | ? | ? | 100 | 97.4 |
| **A8** | M120 | H40 | 102 | 61.7 | 66.7 | 34 | | 34 | ? | 2.94 | 0 | 0 | ? | 2.94 | 62.6 |
| **A9** | M121 | Q11 | 76 | 0 | 2.6 | 74 | | 74 | 50.00 | 100 | 0 | ? | ? | 100 | 97.4 |
| **A10a** | Q11 | DD16a | 206 | 53.3 | 54.9 | 93 | | 93 | 84.95 | 90.33 | 0 | ? | ? | 90.3 | 94.0 |
| **A10b** | Q11 | DD16b | 198 | 54.4 | 57.1 | 85 | | 85 | 77.65 | 88.00 | 0 | ? | ? | 88 | 92.1 |
| **A11** | Q11 | S87 | 187 | 42.9 | 44.9 | 103 | 103 | 0 | | | | | | | |
| **A12a** | T05 | S87a | 113 | 74.5 | 80.5 | 22 | | 22 | 18.18 | 31.82 | 0 | 0 | ? | 31.8 | 80.7 |
| **A12b** | T05 | S87b | 118 | 73.4 | 78.8 | 25 | | 25 | 16.00 | 32.00 | 0 | 0 | ? | 32.0 | 80.2 |
| **A13** | T05 | K169 | 126 | 97.8 | 100 | 0 | | 0 | | | | | | 0 | 97.8 |
| **A14** | L59 | G25 | 130 | 59.1 | 61.5 | 50 | | 50 | 66.00 | 0 | 0 | 56.00 | ? | 66.0 | 84.5 |

**Table S2**, *Contd…*

| Plot No. | Pollen Parent | Ovary Parent | F1 Panicle Density | Hybrid sterility % | Total Grain Sterility % | No. of Fertil e Seeds | F1 Seeds Lost to Pest | No. of Viable F1 Offspring | Non-OP Characters in F1 (%) | | | | | % Viable Progeny with Non-OP traits * | CPF # |
|---|---|---|---|---|---|---|---|---|---|---|---|---|---|---|---|
| | | | | | | | | | Flag Leaf Angle | Awn | Hull Colour | Bran Colour | Notched Belly | | |
| **B1** | AA19 | B103 | 139 | 57.9 | 61.2 | 54 | | 54 | 0.0 | 100.0 | 0 | ? | ? | 100 | 96.7 |
| **B2** | B103 | AA19 | 112 | 98.0 | 100 | 0 | | 0 | | | | | | 0 | 98.0 |
| **B3** | M94 | H01 | 136 | 44.3 | 51.5 | 66 | | 66 | 36.4 | 2.0 | 54.55 | ? | ? | 54.5 | 70.7 |
| **B4** | H01 | M94 | 115 | 97.9 | 100 | 0 | | 0 | | | | | | 0 | 97.9 |
| **B5** | N01 | S72 | 122 | 31.5 | 35.2 | 79 | | 79 | 79.75 | 3.83 | ? | 67.09 | ? | 79.75 | 83.2 |
| **B6** | S72 | N01 | 119 | 66.1 | 75.6 | 29 | | 29 | 31.03 | 93.10 | ? | 0 | 0 | 93.10 | 88.8 |
| **B7** | M91 | J33 | 126 | 60.7 | 67.5 | 41 | | 41 | 100 | ? | 17.07 | ? | ? | 100 | 93.2 |
| **B8** | J33 | M91 | 178 | 47.1 | 55.1 | 80 | | 80 | 85.72 | 100 | 38.75 | ? | ? | 100 | 92.0 |

? = Character common to both parents, so the presence of the character cannot prove its inheritance from a pollen parent.
* - estimated using equation 1.
### - estimated using equation 2.
